# Experimenting with reproducibility in bioinformatics

**DOI:** 10.1101/143503

**Authors:** Yang-Min Kim, Jean-Baptiste Poline, Guillaume Dumas

**Affiliations:** Institut Pasteur, Human Genetics and Cognitive Functions Unit, Paris, France; CNRS UMR 3571 Genes, Synapses and Cognition, Institut Pasteur, Paris, France; University Paris Diderot, Sorbonne Paris Cité, Paris, France; Centre de Bioinformatique, Biostatistique et Biologie Intégrative (C3BI, USR 3756 Institut Pasteur and CNRS), Paris, France; Henry H. Wheeler Jr. Brain Imaging Center, Helen Wills Neuroscience Institute, University of California, Berkeley, California, USA

## Abstract

Reproducibility has been shown to be limited in many scientific fields. This question is a fundamental tenet of the scientific activity, but the related issues of reusability of scientific data are poorly documented. Here, we present a case study of our attempt to reproduce a promising bioinformatics method [1] and illustrate the challenges to use a published method for which code and data were available. First, we tried to re-run the analysis with the code and data provided by the authors. Second, we reimplemented the method in Python to avoid dependency on a MATLAB licence and ease the execution of the code on HPCC (High-Performance Computing Cluster). Third, we assessed reusability of our reimplementation and the quality of our documentation. Then, we experimented with our own software and tested how easy it would be to start from our implementation to reproduce the results, hence attempting to estimate the robustness of the reproducibility. Finally, in a second part, we propose solutions from this case study and other observations to improve reproducibility and research efficiency at the individual and collective level.

**Availability:** last version of StratiPy (Python) with two examples of reproducibility are available at GitHub [2].

## 1 Background

The collective endeavour of science depends on researchers being able to replicate the work of others. In a recent survey of 1,576 researchers, 70% of them admitted having difficulty in reproducing experiments proposed by other scientists [3]. For 50%, this reproducibility issue even concerns with their own experiments. Despite the growing attention on the replication crisis in science [4,5], this controversial subject is far from being new: already in the 17th century, scientists criticized the air pump invented by physicist Robert Boyle because it was too complicated and expensive to build [6].

Several concepts for reproducibility in computational science are closely associated [7,8]. Here we define them as mentioned by K. Whitaker [8]: obtaining the same results using same data and same code is ***Reproducibility***; if code is different, it is ***Robustness***. If we used different data but with the same code, it is ***Replicability***. lastly, using different data and different code is referred as ***Generalisability***. Here we will primarily elaborate on ***Reproducibility*** and ***Robustness***. Indeed, it takes great efforts and competence to overcome all the obstacles on the path to reproduction. The process is costly in resources, both in time and funding. In computational science, there are also many technical barriers ranging from unavailable data to hardware infrastructure [9]. Even when authors provide data and code, the outcome can vary either marginally or fundamentally [10]. Tackling irreproducibility in bioinformatics thus requires considerable effort beyond code and data availability. In most cases, there is a significant gap between apparent executable work (Fig 1 - i.e. above water portion of iceberg) and necessary effort in practice (Fig 1 - i.e. full iceberg). Such effort is nevertheless necessary to increase the consistency of the literature and efficiency of the scientific research process. Indeed, behind reproducibility hides reusability.

**Figure 1:**
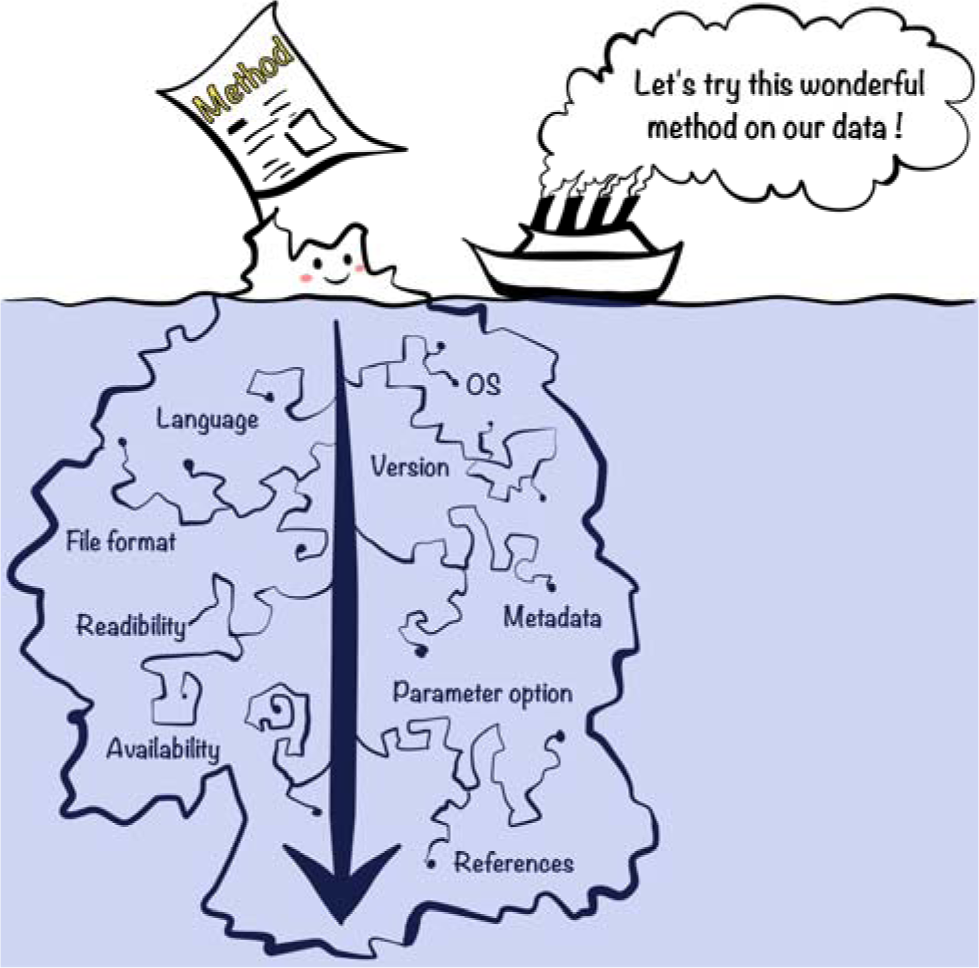
Hidden reproducibility issues like underwater iceberg. Scientific journals readers have the impression that they can almost see the full work of method. But in reality, articles do not take into account adjustment and configuration for significant replication in most cases. Therefore, there is a significant gap between apparent executable work (i.e. above water portion of iceberg) and necessary effort in practice (i.e. full iceberg).

## 2 Reproducibility and Robustness in bioinformatics: a case study

### 2.1 Reproducibility: from MATLAB to MATLAB, OS and environment

Our team studies Autism Spectrum Disorders (ASD), a group of neuro-developmental disorders well known for its heterogeneity. One of the current challenges of our research is to uncover homogeneous subgroups of patients (i.e. stratification) with more precise clinical outcomes, improving their prognosis and treatment [11,12]. An interesting stratification method was recently proposed in the field of cancer research [1], where the authors proposed to combine genetic profiles of patients’ tumours with protein-protein interaction networks to uncover meaningful homogeneous subgroups, a method called Network Based Stratification (NBS).

Before using this NBS method on our data, we studied the method by reproducing results from the original study. We are very grateful to the main authors who kindly provided online all the related data and code, and gave us invaluable input upon request. The authors of this study thus should not be blamed for the difficulty that we experienced in attempting to reproduce and to make more robust their study, as they did more to help reproduce their results than is generally done. Despite their help we experienced a number of difficulties that we document here, hoping that this report will help future researchers to improve the reproducibility of results and reusability of research products.

The first step of our project was to execute the original method code with the given data: reproducibility. The programming code was written in MATLAB, an interpreted language originally developed for linear algebra computations which is easier and faster to write as well as more readable than compiled language such as C, making our reproducibility attempt easier. To improve execution speed, the original authors used a library for MATLAB using executable compiled code MEX file [13] callable from MATLAB: MTIMESX [14], a library with compiled code allowing acceleration of large matrix multiplication. MEX files however are specific to the architecture and have to be recompiled for each Operating System (OS). The original MEX file was initially developed for Linux. Since our lab was using Mac OS X Sierra, the compilation of this MEX file into a mac64 binary required a new version of MTIMESX. It was also necessary to install and to configure properly OpenMP [15], a development library for parallel computing. After this, the original MATLAB code was successfully run in our environment.

These issues are classic, but may not be overcome by researchers with little experience in compilation or installation issues. For these reasons alone, many individuals may turn down the opportunity of reusing code. The next part will focus on code re-implementation, a procedure, which can help understanding the method, but can be even more costly

### 2.2 Robustness: from MATLAB to Python, language and organization

To fully master the method, adapt it to our data, and ease its reuse, we developed a complete open source toolkit of genomic stratification in Python [2]. Python is also an interpreted programming language, but contrary to MATLAB is free of use and has a GPL-compatible license [16]. This is particularly interesting for both robustness and generalizability. Recoding in another language in a different environment will lead to be some unavoidable problems such as variation in low level libraries (e.g. glibc): it is likely that the outcomes will vary even if the same algorithm is implemented. In addition, we rely on Python packages to perform visualization or linear algebra computations (e.g. Matplotlib, SciPy, NumPy), and results may depend on these packages versions. Python is currently in a transitional period between two major versions 2 and 3. We chose to write the code in Python 3, which is the current recommendation.

### 2.2.1 Metadata and file formats

Even if the original code could be run, we had to handle several file formats to check and understand the structure of the original data. For instance the data on patients with cancer data was provided by The Cancer Genome (TCGA) [17] and made available in a MATLAB *.mat* file format. Thanks to SciPy, Python can load all versions through v7.2 MATLAB files. To read v7.3 *.mat* files, we however needed an HDF5 Python library. Moreover, the original authors had denoted download dates of patients’ data of TCGA, thereby clarifying source of data. But in the absence of structural metadata, it was not always obvious how to interpret patients’ dataset variables (e.g. patient ID, gene ID, phenotype). Fig 2 shows an analogy between robustness issues and road transport: driving in a different environment (e.g. OS), we attempt to obtain identical results (i.e. to reach the same location) using the same input data of TCGA (i.e. gasoline). But there is a difficult to transfer proposed method (i.e. engine) from one programming language to another (i.e. MATLAB and Python roads).

**Figure 2:**
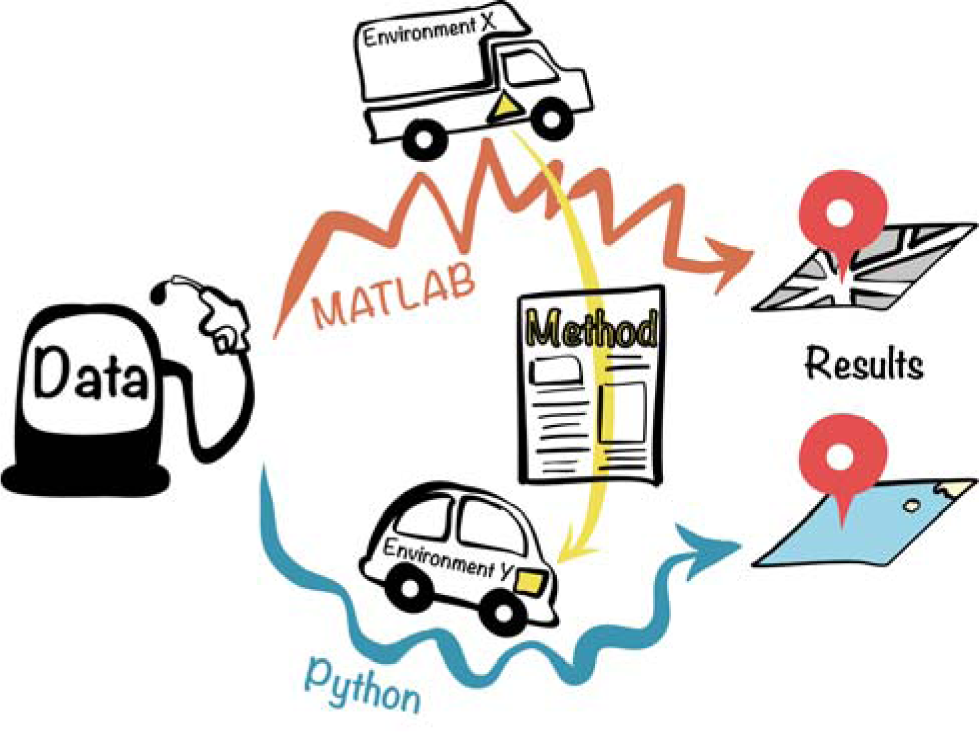
Analogy between robustness issues and road transport. The aim is to achieve same output (i.e. to reach the same location) using published methods (i.e. engine). Despite the same input data (i.e. gasoline), we obtained different results due to different programming languages —e.g. MATLAB and Python— (i.e. different roadways) and environments (i.e. different vehicles).

### 2.2.2 Codes and parameters

Once the environment and file format issues were resolved, the code was finally executable with genetic data. Unfortunately, several attempts produced error messages. Alternatively, “unexpected” results were obtained: e.g. during the application of hierarchical clustering, we used the clustering tools of SciPy. Both SciPy and MATLAB (MathWorks) functions offer seven linkage methods, however, SciPy’s default option (single method) [18] differs from MATLAB’s default option (UPGMA or average method) [19], which was used in the original study. Another key example is the value of one of the most important parameters of the method, the graph regulator factor, which was not clarified in the original paper. We believed that this factor had a constant value of 1.0 until we found in the code that during iterations, its value was changing and converged to a high optimal value (~1800). Therefore, we obtained different results from the original NBS at the beginning (Fig 3). We observed heterogeneous subgroups instead of obtaining homogeneous clusters. No or little explanation on the parameter choices can explain variability in the results as we explored the possible parameters. Moreover, during our attempts to run the original code to understand the causes of the errors, we realized that some parts of the code were not run anymore (e.g. discarded work, remaining traces of debugging) which made the attempt to understand the implementation harder.

**Figure 3:**
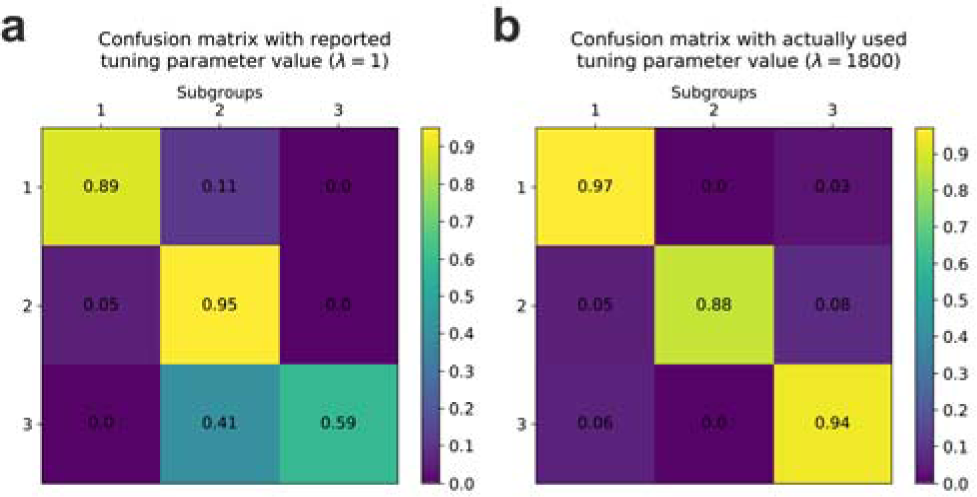
Normalized confusion matrices between original and replicated results. Before (a) and after (b) applying appropriate value of graph regularization factor on NBS method. Each row or column corresponds to a subgroup of patients (here three sub-groups). The diagonal elements show the frequency of correct classifications for each subgroup: a high value indicates a correct prediction.

Readers are proposed to reproduce those two confusion matrices (Fig 3) using different practical tools: GitHub, Docker and Jupyter/IPython notebook, which are further described below.

### 2.2.3 Jupyter/IPython

During the re-cording process, we used an enhanced Python interpreter to debug: IPython, an interactive shell supporting both Python 2 and 3. Since the dataset is large and the execution takes a significant amount of time, we used IPython to re-run interactively some sub-sections of the script, which is one of the most helpful features. IPython can be integrated in the web interface Jupyter Notebook, offering an advanced structure for mixing code and documentation. For instance, verifying intermediate results by plotting helped us to better understand the original code. While the Jupyter/IPython notebook was therefore initially convenient, it does not scale well and is not well adapted to versioning. However, ability of mixing code with document text is very useful for tutorials: a user of the code can read documentation (docstring), text explanations, and see how to run the code, explore parameters and visualize results in the browser. Our work on NBS, as related here, can be reproduced with a Jupyter/IPython notebook available on our GitHub [2]. You can find more examples and several helpful links on this “gallery of interesting Jupyter Notebooks” [20], which even contains a section about “Reproducible academic publications”.

### 2.3 Reproducibility of Robustness: from Python to Python

Besides Jupyter/IPython notebook, we used versioning tools like the git code version control system (VCS) to document the development of our Python code. Git is arguably one of the most powerful VCS, allowing easy development of branches and helping us to work together as a distributed team (Paris, Berkeley) on the same project. This project, *StratiPy*, is hosted on GitHub, a web-based Git repository hosting service [2]. While the original code was not available on GitHub, the main authors shared their code on a website. This should be sufficient for our purpose, but makes it less easy to collaborate on code. While working on our GitHub repository, several researchers from all over the world contacted us about our robustness experiment. Not only GitHub supports a better organization of projects, it also facilitates the collaboration of open-source software projects, thanks to several social network functions [21]. We tried to comply with open source coding standards and to learn how to efficiently use Git and GitHub. Both required considerable efforts on the short-term but brought clear benefits on the long-term, especially regarding collaboration and debugging.

We then attempted to re-run and reproduce the results we obtained on another platform. While the Python code was developed under Mac OS X Sierra (10.12) we used an Ubuntu 16.04.1 (Xenial) computer to test the Python implementation. Some additional issues emerged. First, our initial documentation was not complete enough to know which packages were required and how to launch the code. Second, the code was very slow to the extent that it was impractical to run it on a laptop because the Numpy package had not been compiled with BLAS (Basic Linear Algebra Subprograms), low-level routines performing basic vector and matrix operations. Last, there was (initially) no easy way to check whether the results obtained on a different architecture were the expected ones. We added documentation and tests on the results files md5sum to solve this. To summarize, although the reuse and reproducibility of the results of the developed package were improved, these were far from being optimal.

## 3 Potential solutions: from local to global

### 3.1 Act locally: simple practices and available tools

Given the observed difficulties, we draw some conclusions on this reproducibility case study experiment and suggest some practices and tools. In addition to this guidance, computer scientists are strongly encouraged to follow detailed advice of Wilson et al. such as modularizing and re-using code, unite testing, document design, data management, and project organization [4,22]. Sandve and colleagues [23] also suggest to keep the data provenance with recording all intermediate results.

### 3.1.1 Environment

Container technologies such as Docker [24], Vagrant [25], or Singularity [26,27] (easily works in cluster environments) are becoming a standard solution to installation issues. These rely however on competencies that we think few biologists possess today. Also, while the container will encapsulate everything needed for the software execution, it could be hard to develop in a container. For instance, running Jupyter/IPython notebooks in Docker’s container requires certain knowledge of computer science (e.g. advanced port forwarding), which can become a discouraging task. Therefore, we decided to propose two options in our example implementation of reproducibility: 1) a step-by-step process to follow in a Jupyter/IPython notebook; or 2) a Docker container ready to be built and run. Nevertheless, mastering Docker –or other container tools– will become an important skill for computational reproducible researchers.

### 3.1.2 Metadata

Standard metadata are vital for an efficient documentation of both data and software. In our example, we still lack the standard lexicon to document the data as well as documenting the software: e.g. using HDF5 file instead of *.mat* file is more suitable to store patient's’ data. We however aim to follow the recommendations by Stodden *et al.* [28]: “*Software metadata should include, at a minimum, the title, authors, version, language, license, Uniform Resource Identifier/DOI, software description (including purpose, inputs, outputs, dependencies), and execution requirements”.* The more comprehensive is the metadata description, the more likely the reuse will be both efficient and appropriate [29].

### 3.1.3 Write readable code

Anyone who has spent time to understand someone else’s code would advise some simple basic rules to help make the code readable and understandable.

First, the structure of the program should be clear and easily accessible. Second, good concise code documentation and naming convention will help readability. Third, the code should not contain left-overs of previously tested solutions. When a solution takes a long time to compute, an option to store it locally can be proposed. Using standard coding and documentation conventions (e.g. PEP 8 and PEP 257 in Python [30,31]) with detailed comments and references of papers makes the code more accessible. When an algorithm from another paper is used, any modification should be explained and discussed in the paper as well as in the code. All these remarks are not necessarily obvious especially if the developer is working on her/his own, and to some extent “writes for her/himself”. We advocate for researchers to write code “for their colleagues”, hence, the opinion and notice of co-working or partner laboratories should be very helpful. Furthermore, the collaboration between researchers working on different environments can more easily isolate reproducibility problems. In the future, journals may consider review of code as part of the standard review process.

### 3.1.4 Test the code

To check if the code is yielding a correct answer, software developers associate test suites (unit tests or integration tests) with their software. While we developed only a few tests in this project, we realize that this has a number of advantages, such as checking if the software installation seems correct, check if updates in the operating system impact the results, etc. This does not in general validate the method, but at least provides a basic check. In our case, we propose to check for the integrity of the data and for the results of some key processing.

## 3.2 Think globally: from education to community standards

### 3.2.1 Training the new generation of scientists to digital tools and practices

Unlike theoretical and academic courses and projects, software testing systems are well developed in industry since software quality is not the priority in Academia [32]. For a student, discovering and learning this core system of reproducibility, possibly during an internship in cooperation with industry, is a great opportunity for her/his future. Furthermore, as Internet applications in science are growing, networks of scientists and developers are forming and provide learning opportunities on the development practices. For instance, software developers have recently adopted “agile” practices and fast prototyping, test based development, etc. Some of these ideas and practices can —and should— be adapted to scientific software development.

The training in coding is still too limited for biologists. Often, it is self-training, from searching answers on Stack Overflow or equivalent. Despite efforts by organizations such as software [33] or data carpentry [34] and the growing demand for ‘data scientists’ in life science, university training on coding practices is still not enough generalized. The difficulty to access and understand code may lead to applying code blindly without checking the validity of the results: often, scientists may prefer to believe that the results are correct because of the time that would be needed to check the validity of the results. Mastering a package such that results are truly understood can take a long time, as it was the case in our experiment.

Academia could instruct young scientists best practices for reproducibility. For instance, Hothorn and Leisch organized a reproducibility workshop gathering mostly PhD students and young postdocs specialized in bioinformatics and biostatistics. Then they evaluated 100 random sample papers from *Bioinformatics* [5]. Their study revealed how such a workshop can raise young scientists awareness about “*what makes reproduction easy or hard at first hand”*. Indeed, they found out that only a third of the original papers and two-thirds for applications notes had given access to the source code of software used.

### 3.2.2 Standard consensus dataset and workflow system

We propose here that bioinformatics methods publications are systematically accompanied with a test dataset, code source and some basic tests. As the method is tested on new datasets, the number of tests of the method would increase in number and cover a wider range of applications. We give a first example with our NBS re-implementation. We develop below how this could generalize and what would be the benefit for the scientific community. In a sense, we propose to use the software development test framework idea but apply it to the scientific context.

A schematic overview of workflow system is shown in Fig 4. The core of this system would be a standard consensus dataset used to validate methods. For instance in the field of machine learning, standard image databases are widely used for training and testing (e.g. MNIST for handwritten digits [35]). In the case of our proposal, data could be classified in general categories such as binary, text, image (A, B, C in Fig 4 b), and with sub-categories to introduce criteria such as size, quantitative/qualitative, discrete/continuous using a tagging system (e.g. A-2, B-1, C-5 in Fig 4 b). Dataset could be issued from simulations or from acquisition, and would validate a method on a particular component. This workflow system will help scientists that cannot release their data because of privacy issues (Fig 4 a.1) (although these can often be overcome) but also give access to data and tests to a wide community. Scientists can use only standard data at the beginning of the project. And if there is no appropriate data, they have to suggest a new standard data.

**Figure 4:**
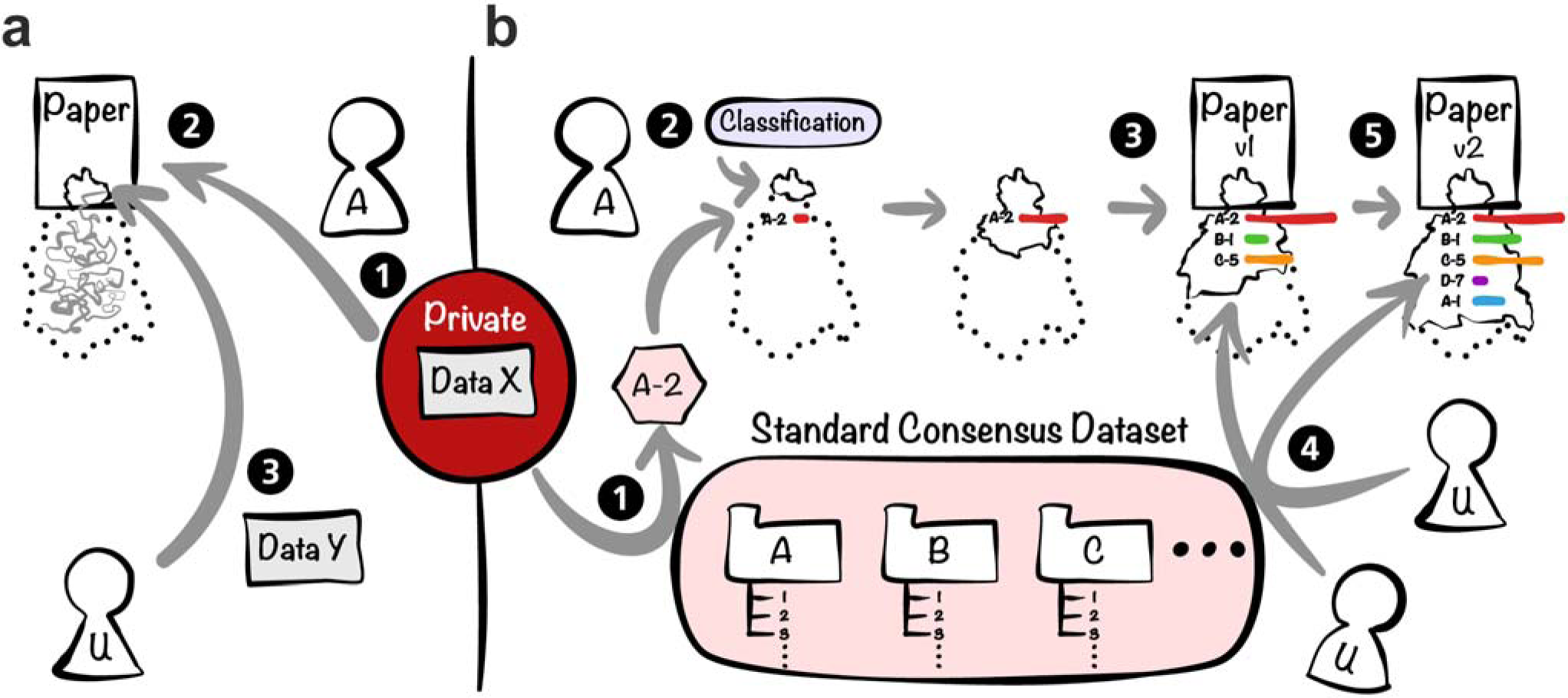
Working principles of Workflow system with private data. Figure 4a shows a classical workflow: (a.1) Authors (“A”) take private data; (a.2) Authors publish their method and corresponding outputs/results; (a.3) Users (“U”) having their own data find a relevant paper but will be lost in the labyrinth of reproducibility. Figure 4b shows workflow with standard consensus dataset: (b.1) If authors work with their own data, they must identify corresponding standard data tag(s) (e.g. A-2); (b.2) Users choose a method category (e.g. "Classification"); (b.3) Reproducibility profile with standard data is progressively built with method upgrade throughout publication of the first version. Color corresponds to method category depending score and bar length corresponds to progression of replication test; (b.4) Users can test proposed method with other data standards and thus participate to enhancement of the reproducibility profile; (b.5) Thanks to the collective work on testing, the method could be optimized and authors can upgrade their initial paper (versioning).

Roughly, we divide those who interact with scientific software or analysis code in two categories. First, the authors (“A”) who propose a method and need to verify its validity and usefulness with public and/or their own – often private – data. Second, the users (“U”, e.g. developers, engineers, bioinformaticians) who need to test and evaluate the proposed methods with other data.

When authors launch a research about a method, this method must belong to a general category of methods (e.g. classification, regression) and could have a reproducibility profile, which will progressively be built by authors and users (Fig 4 b.3, b.4). Even the information of which method does or does not work with a standard data is a crucial information for future work. During optimization of project, the programming code with guide should be accessible although authors do not publish. To achieve current work, they can also post on preprint servers such as bioRxiv [36,37] associated with a GitHub repository by digital object identifiers (DOI).

Furthermore, users who test and approve reproducibility on original or new data could be credited and recognized by the scientific and developer communities (i.e. Stack Overflow, GitHub). This workflow system thus could facilitate the gathering of diverse users of the science community.

## 4 Conclusion and perspective

Across the scientific fields, the reproducibility issue is seen as a growing concern. Before reusing a published method, we attempted to reproduce the initial results and recoded the method to have a deep understanding of it. The investment in time to verify a previously published method can be more important than the work needed to publish a new paper. Despite the willingness of the authors to share their tool and help us in our work, we have faced computational reproducibility and robustness problems due to compatibility between environments, programming languages and software versions, choice of parameters, etc. In addition to individual effort to write well documented and readable code, we recommend to use online repositories and tools to help other scientists in their exploration of the method: Docker for environment standardization, GitHub for code version management, and Jupyter notebooks for demonstration and tutorial [20,21,24]. Scientists are strongly encouraged to adopt such practices, not only for writing code but also manuscripts [4]. At the community level, we should enhance the cooperation between academic education and industry to foster a new generation of well-trained scientists in software development. For instance, Academia-Industry Software Quality & Testing summit (AISTQ) has organised conference in order to encourage collaboration between Academia and Industry [38]. Here, we propose a workflow system where the community uses standard datasets to validate tools. The proposed method success on data profile will be evaluated continuously with new datasets. Eventually, data and software can be versioned and cited to give credit to the individuals who have contributed to these building blocks of Science. This workflow is not merely a reproducibility validation tool, it is an attempt to make research product more reusable by the community using an online platform, beyond the publication process. Such system could be seen as a generalisation of already existing workflow systems such as Galaxy or GATK, integrating data provenance [39,40]. Some top-down initiatives already provide some incentives for such a process i.e. Horizon 2020 (H2020) [41] project of the European Commission (EC) mandates open access of research data, while respecting security and liability. H2020 supports OpenAIRE [42], a technical infrastructure of the open access, which allows the interconnection between projects, publications, datasets, and author information across Europe. Thanks to common guidelines, OpenAIRE interoperates with other web-based generalist scientific data repositories such as Zenodo, hosted by CERN, which allows combining data and GitHub repository using DOI. The Open Science Framework also hosts data and software for a given project [43]. Respecting standard guidelines to transparently communicate the scientific work is a key step towards tackling irreproducibility and insures a robust scientific endeavor.

## Key points

- Main barrier for reproducibility is in the lack of compatibility between environments, programming languages, software versions, etc.
- At the individual level, the key is in research practices such as proper code and data documentation and exploitation of online repositories and collaborative tools.
- At the community level, we propose a workflow system where standard consensus datasets are used to validate new methods and foster their generalizability.

